# A novel sulfatase for acesulfame degradation in wastewater treatment plants as evidenced from *Shinella* strains

**DOI:** 10.1101/2024.03.04.583314

**Authors:** Yu Liu, Thore Rohwerder, Maria L. Bonatelli, Theda von Postel, Sabine Kleinsteuber, Lorenz Adrian, Chang Ding

**Author notes:** To EST correspondence should be addressed: Chang Ding, Helmholtz Centre for Environmental Research – UFZ, Department of Molecular Environmental Biotechnology, Permoserstraße 15, 04318 Leipzig, Germany; Thore Rohwerder, Helmholtz Centre for Environmental Research – UFZ, Department of Microbial Biotechnology, Permoserstraße 15, 04318 Leipzig, Germany.

## Abstract

The artificial sweetener acesulfame is a persistent pollutant in wastewater worldwide. So far, only a few bacterial isolates were recently found to degrade acesulfame efficiently. In *Bosea* and *Chelatococcus* strains, a Mn^2+^-dependent metallo-β-lactamase-type sulfatase and an amidase signature family enzyme catalyze acesulfame hydrolysis via acetoacetamide-N-sulfonate (ANSA) to acetoacetate. Here, we describe a new acesulfame sulfatase in *Shinella* strains isolated from German wastewater treatment plants. Their genomes do not encode the Mn^2+^-dependent sulfatase. Instead, a formylglycine-dependent sulfatase gene was found, together with the ANSA amidase gene on a plasmid shared by all known acesulfame-degrading *Shinella* strains. Heterologous expression, shotgun proteomics and size exclusion chromatography corroborated the physiological function of the *Shinella* enzyme as a Mn^2+^-independent acesulfame sulfatase. Since both the *Bosea*/*Chelatococcus* sulfatase and the novel *Shinella* sulfatase are absent in other bacterial genomes or metagenome assembled genomes, we surveyed 60 tera base pairs of wastewater-associated metagenome raw datasets. The *Bosea*/*Chelatococcus* sulfatase gene was regularly found from 2014 on, particularly in North America, Europe and East Asia, whereas the *Shinella* sulfatase gene was first detected in 2020. The complete *Shinella* pathway is only present in five datasets from China, Finland and Mexico, suggesting that it emerged quite recently in wastewater treatment facilities.

**Synopsis:** A novel sulfatase was identified that hydrolyzes the once recalcitrant xenobiotic acesulfame. Surveying metagenome datasets revealed the recent emergence of gene homologs encoding this sulfatase in wastewater treatment systems worldwide.

## Introduction

Acesulfame is one of the most commonly used artificial sweeteners in low-calorie food and beverages, pharmaceuticals and cosmetics.^1,2^ The global consumption of acesulfame, which was excreted unchanged via the kidney, led to its widespread presence in domestic wastewater (typically between 10 and 100 μg L^-1^),^3^ groundwater (up to 5 μg L^-1^),^4^ seawater (0.57-9.9 μg L^-1^),^5^ and tap water (up to 2.6 μg L^-1^).^6^ Acesulfame concentrations of up to 2.5 mg L^-1^ were reported in effluents from European wastewater treatment plants (WWTPs).^7^ Even though the approved usage of acesulfame as a food additive indicates its low estimated risks for human health and the environment, several studies highlighted potential risks associated with acesulfame and its transformation products. Acesulfame acts as an inhibitor of P-glycoprotein, thereby reducing the detoxification capacity of the liver.^8^ It also induces uterine hypercontraction upon long-term high-dose exposure.^9^ Moreover, it can be transformed into more persistent and more toxic products by chlorination,^10^ UV irradiation,^11^ and TiO_2_-assisted photolysis,^12,13^ posing a long-term potential risk for aquatic environments. For example, the toxicity of acesulfame transformation products after UV/TiO_2_ treatment was found to be magnified by a factor of 575 compared to that of acesulfame itself.^12^ Other evidence indicates that acesulfame might cause DNA damage^14^ through the formation of complexes in the minor groove of DNA.^15^ Recent studies reported oxidative stress in fish^16^ and neurotoxic effects in *Daphnia magna*^17^ upon acesulfame exposure. Additionally, it has been observed that acesulfame can promote the horizontal dissemination of antibiotic resistance genes in pure culture systems, ^18^ in gut microbiome,^19^ and during anaerobic digestion.^20^

Previously, acesulfame was regarded as an ideal tracer to identify anthropogenic contamination sources^6,21^ due to its persistence and mobility in aquatic systems.^22^ However, recent studies showed that acesulfame can actually be biodegraded,^23,24^ which makes its use as conservative tracer obsolete. Nevertheless, since biodegradation is temperature-dependent,^24,25^ acesulfame can still be used as a transient tracer in cold season (<10°C).^26,27^

Thus far, 14 bacterial strains belonging to the genera *Bosea* (strains 3-1B and 100-5),^28^ *Chelatococcus* (strains 1g-2, 1g-11, WSA4-1, WSC3-1, WSD1-1, WSG2-a, YT9 and HY11),^28-30^ and *Shinella* (strains HY16, YE25, YZ44, and WSC3-e)^30,31^ have been isolated from wastewater habitats that were able to grow with acesulfame as sole carbon and energy source. In addition, we recently isolated *Shinella* sp. WSD5-1 which was also able to grow on acesulfame. Among all these strains, *Bosea* sp. 3-1B was the first acesulfame-degrading bacterial isolate worldwide, which was obtained from a German treatment wetland sample taken in 2015.^28^ In *Bosea* and *Chelatococcus* strains, acesulfame is hydrolyzed via acetoacetamide-N-sulfonate (ANSA) to acetoacetate and sulfamate. In this pathway, a Mn^2+^-dependent metallo-β-lactamase (MBL)-type sulfatase catalyzes the nucleophilic attack of acesulfame to form ANSA.^30^ Subsequently, the amide bond of ANSA is cleaved by an amidase signature family enzyme.^30^ However, in the genomes of the above-mentioned *Shinella* strains^30,31^ and also in our newly sequenced strain *Shinella* sp. WSD5-1, the gene for the Mn^2+^-dependent MBL-type sulfatase is missing, while the ANSA amidase gene known from the *Bosea*/*Chelatococcus* genomes is conserved.

Therefore, we hypothesized that in *Shinella* strains a different hydrolase may be responsible for the acesulfame hydrolysis step and aimed to identify this novel acesulfame-hydrolyzing enzyme in *Shinella* sp. WSC3-e. Through heterologous expression of a candidate gene, activity assays and proteomic analysis, we discovered a novel formylglycine-dependent sulfatase and experimentally confirmed its acesulfame-hydrolyzing activity. To obtain indications about the historic occurrence and geographic distribution of the acesulfame degradation gene sequences, we analyzed wastewater-derived metagenome raw sequence datasets for the occurrence of both the *Shinella* and *Bosea*/*Chelatococcus* acesulfame sulfatase and ANSA amidase genes.

## Material and Methods

### Cultivation of *Shinella* strains

*Shinella* sp. WSC3-e (genome assembly European Nucleotide Archive, ENA, and GenBank ID: GCA_945994535.2) was previously isolated from activated sludge of the WWTP Rosental Leipzig, Germany.^30^ *Shinella* sp. WSD5-1 (ENA/GenBank ID: GCA_963942435.1) was isolated from the WWTP of Markkleeberg, Germany, which was sampled in 2020. Both strains showed the capability of growing on acesulfame as the sole carbon and energy source. To study growth and degradation kinetics, strain WSC3-e was cultivated in Brunner mineral medium (DSMZ 462, containing 0.15 µM MnCl_2_) with 30 mM acesulfame at 30°C. A Mn^2+^-rich control was incubated in the same medium supplemented with 3.7 μM MnCl_2_. For DNA extraction, *Shinella* sp. WSD5-1 was grown on 5 mM acesulfame in mineral salt medium (DSMZ 461) at 30°C.

### Genome sequencing of strain WSD5-1

DNA of acesulfame-grown cells was extracted with MagAttract HMW DNA Kit (QIAGEN, Hilden, Germany), according to the manufacturer’s instruction. DNA integrity was checked by gel electrophoresis, while DNA concentration was measured with the Qubit dsDNA BR Assay Kit (Thermo Fisher Scientific, Rockford, IL, USA). The genome was sequenced with both long- and short-read techniques. Long-read sequencing was performed with MinION Mk1B platform (Oxford Nanopore Technologies, Oxford, United Kingdom). Library was prepared with the ligation sequencing kit V14 (SQK-LSK114), and sequencing was done using the flow cell R10.4.1 with the software MinKNOW UI v.23.07.15. Basecalling was performed with the software dorado v.0.4.2 (Oxford Nanopore Technologies). Filtlong v.0.2.1 (github.com/rrwick/Filtlong) was used to filter out the 40% worst reads. Short-read sequencing was done on the Illumina NovaSeq platform (2 × 150 bp; Azenta, Leipzig, Germany). Assembly was conducted with flye v.2.9.2,^32^and genome polishing was done with pilon v.1.23.^33^ All software tools were used with default parameters. Genome annotation, classification and quality measurements were done as previously described.^32^ *Shinella* sp. WSD5-1 genome was deposited on ENA under the project PRJEB50809.

### Comparison of *Shinella* genomes

The FastANI software^34^ in the Proksee system^35^ was used to calculate the average nucleotide identity of the *Shinella* sp. WSC3-e genome with the genomes of strains *Shinella* sp. WSD5-1, YE25 (GenBank ID: GCA_028534295.1) and HY16 (GenBank ID: GCA_028534175.1). Proksee was also used for the alignment of the YE25 contigs against the WSC3-e genome.

### Heterologous expression of the sulfatase gene SHIWSC3_PJ0001

SHIWSC3_PJ0001 was synthesized and cloned (via *Nde*I/*Bam*HI sites) by GenScript (Oxford, United Kingdom) into the expression vector pET-28a (+)-TEV (using the gene without the initial 36 bp). The recombinant plasmid was transformed into *E. coli* Lemo21 (DE3) (New England Biolabs), which was grown at 30°C in lysogeny broth with 30 mg L^-1^ chloramphenicol and 50 mg L^-1^ kanamycin until an optical density (600 nm) of 0.5 was reached. Then, 0.4 mM of isopropyl β-d-1-thiogalactopyranoside were added and incubated further at 30°C for 4 h. Cells were harvested by centrifugation. As a control experiment, a gene encoding a 465-aa amidase signature enzyme from *Bosea* sp. 100-5 (BOSEA1005_40016), which is not involved in acesulfame or ANSA degradation,^30^ was expressed from pET-28a (+)-TEV under the same conditions.

### Acesulfame hydrolyzing activity in *E. coli* crude extracts

Protein extracts prepared in lysis buffer (50 mM Tris-HCl with 2 mM dithiothreitol, DTT, 5 mM MgCl_2_ and 10% glycerol, pH 7.8) were disrupted in a mixer mill with glass beads.^30^ Crude extracts were obtained by taking supernatant after centrifugation at 20,000 x g and 4°C for 20 min. These were tested for acesulfame hydrolyzing activity. The activity assay contained 50 mM Tris-HCl (pH 7.8), 10% glycerol, 2.25 mM acesulfame (potassium salt, 99% purity, Merck, Darmstadt, Germany), 2 mM adenosine 5’-triphosphate (ATP), 2 mM DTT, 5 mM MgCl_2_ and crude extract (added in the end to start the reaction). Incubation was done at 30°C and 300 rpm. Reactions were stopped by adding two volumes of 10 mM sodium malonate buffer (pH 4.0, preheated to 60°C) to assay samples and incubating the mixture at 60°C for 5 min. Then, acesulfame and ANSA concentrations were quantified.

### Size exclusion chromatography (SEC)

To obtain protein fractions from SEC with acesulfame hydrolyzing activity, 200 µL crude extract from *Shinella* sp. WSC3-e cells (1.6 µg protein µL^-1^) was loaded onto a Superdex 200 10/300 column (GE Healthcare, Chicago, USA), which was equilibrated with a buffer containing 50 mM Tris-HCl (pH 7.84), 10% glycerol, 2 mM ATP, 2 mM DTT and 5 mM MgCl_2_ at 0.5 mL min^-1^. Fractions of 0.5 mL between elution volumes 7 mL and 22 mL were collected, which were used for both acesulfame hydrolyzing activity assay and proteomic analysis. The activity assay contained 78 µL SEC fraction subsample and 2 µL of 90 mM acesulfame stock solution (final concentration: 2.25 mM) and was incubated at 30°C and 400 rpm for 40 min before being stopped as described above. Hydrolase activity was calculated from end-point acesulfame concentrations. For assigning SEC retention times to molecular masses, the system was calibrated using the Protein Standard Mix 15-600 kDa (Sigma-Aldrich, Darmstadt, Germany). Calibration was done under a flow rate of 0.5 mL min^-1^ with a mobile phase containing 50 mM Tris-HCl buffer (pH 7.5), 10% glycerol, and 0.03% (w/v) digitonin.

### Shotgun proteomics

*Shinella* sp. WSC3-e crude extract and SEC fractions were analyzed using shotgun proteomics. Each SEC fraction subsample (70 µL) was spiked with 4 µL glyceraldehyde-3-phosphate dehydrogenase (GAPDH, from *Staphylococcus aureus* MRSA252, NCBI ID: WP_000279414, 336 aa residues) as the internal standard to account for sample to sample variation among fractions. Purified GAPDH (concentration between 16 and 20 mg mL^-1^) was diluted 1,000 times in water before usage. Crude cell extract (120 µL, 0.33 µg protein µL^-1^) was processed directly in the following steps without adding GAPDH.

An equal volume of sodium deoxycholate 10% (w/v) was added to all samples in order to denature proteins. Proteins were reduced with 12 mM DTT in 45 min at 37°C and then alkylated with 40 mM iodoacetamide in 45 min at room temperature (both were final concentrations). Before trypsin digestion, samples were diluted with 100 mM ammonium bicarbonate to a final concentration of 1% (w/v) sodium deoxycholate. Trypsin digestion was done by adding 5 µL reductively methylated trypsin (Promega, Madison, USA) to each sample and incubating the samples overnight at 37°C. Digestion was stopped by adding neat formic acid to a final concentration of 2% (v/v). Precipitated material was carefully removed by centrifugation twice at 16,000 x g and 4°C for 10 min. Digested samples were desalted by using 100-µL C_18_ ziptips (Pierce^TM^, Thermo Fisher Scientific, Massachusetts, USA) and measured on a nano-LC-MS/MS (Orbitrap) as described previously.^36^

Protein identification was conducted using the Proteome Discoverer 2.4 (Thermo Fisher Scientific, Massachusetts, USA) applying the SequestHT search engine with the protein database of *Shinella* sp. WSC3-e. Other parameter settings were as described previously^36^ except that mass tolerance for fragment ion was reduced from 0.5 Da to 0.1 Da. Protein and peptide abundance values were calculated by intensity-based label free quantification using the Minora node implemented in Proteome Discoverer.^36^ Protein quantities were normalized using the GAPDH intensity in each SEC sample. Hierarchical cluster analysis was done in OriginPro, Version 2023 (OriginLab Corporation, Northampton, Massachusetts, USA). Correlation was applied to cluster variables into five clusters. Group average was set as the linkage method to calculate the distance among clusters. Clustroid was found by using the sum of distances measured from all other variables in the cluster. The hierarchical tree was shown in a dendrogram plot. The proteomics data were deposited to the ProteomeXchange via the PRIDE partner repository with the number PXD046105.

### Analytical methods

Protein concentration was determined with the bicinchoninic acid (BCA) method, using the Pierce BCA Protein Assay Kit (Thermo Fisher Scientific, Massachusetts, USA) according to the manufacturer’s instruction or Bradford reagent (AppliChem, Darmstadt, Germany), using bovine serum albumin as the standard. Acesulfame and ANSA were quantified using HPLC and LC-MS/MS, respectively, as previously described.^30^

### Database search

The NCBI blastp tool was used to search for sequences related to the acesulfame sulfatase (SHIWSC3_PJ0001 gene product) and ANSA amidase (SHIWSC3_PJ0040 gene product) from *Shinella* sp. WSC3-e against the NCBI non-redundant protein sequence database (February 2024), which covers all annotated genomes and metagenomes. Additionally, other whole genome shotgun sequencing and metatranscriptome datasets in the NCBI and JGI databases (February 2024) originating from wastewater, activated sludge or bioreactor samples were searched with blastn (megablast, NCBI default settings, without filters and masks) for sequences similar to the *Shinella* acesulfame sulfatase and ANSA amidase genes. Furthermore, raw sequence files of shotgun metagenome and metatranscriptome projects in the context of wastewater environments were downloaded from the NCBI Sequence Read Archive (SRA) (Table S1) and queried with blastn (megablast, NCBI default settings, without filters and masks) at the high-performance-computing cluster EVE (UFZ/iDiv) using the acesulfame sulfatase and ANSA amidase genes from *Bosea*/*Chelatococcus*^30^ and *Shinella* as query. A significant match was defined as the alignment of single ≥150-bp reads or the combination of shorter sequences (e.g., two 100-bp reads) resulting in ≥140 bp query coverage at ≥97% identity. Reads having incomplete alignment due to matching only with gene start or end regions were also checked for possible alignments with sequences flanking the corresponding sulfatase and amidase genes in the respective genomes (*Shinella* sp. WSC3-e and *Bosea* sp. 100-5). If those reads match the flanking sequences and fulfill the alignment criteria above, they were also considered as significant matches. A sampling site containing reads of significant matches to a query gene at a specific year was counted as one positive sampling site of this query gene, regardless how many reads are associated with the detection.

## Results

### Genomes of acesulfame-degrading *Shinella* strains

The previously sequenced genomes of acesulfame-degrading *Shinella* strains YE25, HY16^31^ and WSC3-e^30^ as well as the genome of the new isolate *Shinella* sp. WSD5-1 were compared. Although isolated from different locations, strains YE25 (WWTP Sha Tin, Hong Kong, China), WSD5-1 (WWTP, Markkleeberg, Germany) and WSC3-e (WWTP Rosental, Leipzig, Germany) possess almost identical genomes (≥99% sequence identity). Moreover, replicon organization is similar in these strains, representing a chromosome of 4.7 mega base pairs (Mbp) and up to 12 plasmids (Table S2, Figure S1). In two cases, two smaller plasmids of strain YE25 and WSD5-1 were rearranged to larger ones in strain WSC3-e (Figure S1). All three genomes hold a total sequence length of 7.8 Mbp. In contrast, the genome of HY16 (also from WWTP Sha Tin) only amounts to 7.3 Mbp and shares about 74% of the sequence (at ≥90.8% identity) with the other three *Shinella* strains.

### One of the plasmids harbors genes for acesulfame degradation

A closer inspection of the genomes revealed that the four *Shinella* strains have an almost identical 41.6 kilo base-pair (kbp) plasmid (plasmid PJ in strain WSC3-e, ENA/GenBank ID: OZ000540) with 100% conservation of all coding sequences (termed *Shinella* <u>a</u>cesulfame <u>u</u>tilization <u>p</u>athway, AUP, plasmid) (Figure 1). The *Shinella* AUP plasmid harbors the ANSA amidase gene cluster encoding the second step of the acesulfame degradation pathway previously described for *Bosea* and *Chelatococcus* strains.^30^ Among all the AUP-bearing strains, the ANSA amidase gene cluster shows variation in length, ranging from two coding sequences in *Bosea* sp. 100-5 to four in *Chelatococcus* sp. WSC3-1 and 1g-11.^30^ The *Shinella* ANSA amidase gene cluster shows the highest similarity with the three coding sequences-bearing cluster found in strain *Bosea* sp. 3-1B (Figure 1). Besides the gene for the ANSA amidase (locus tag SHIWSC3_PJ0040), two other coding sequences (SHIWSC3_0052 and _0041) are present in the *Shinella* ANSA amidase gene cluster, encoding a TauE/SafE export protein and a LysR-like transcriptional regulator, respectively. In addition to the ANSA amidase gene cluster, the *Shinella* AUP plasmid encodes several backbone genes involved in conjugative transfer, replication and maintenance, which are homologous to and syntenic with their counterparts on the *Bosea* AUP plasmid (Figure 1), e.g., the transfer and replication genes *traA*, *traG* and *parA*. This is indicative of a common origin of the plasmids.

**Figure 1.**
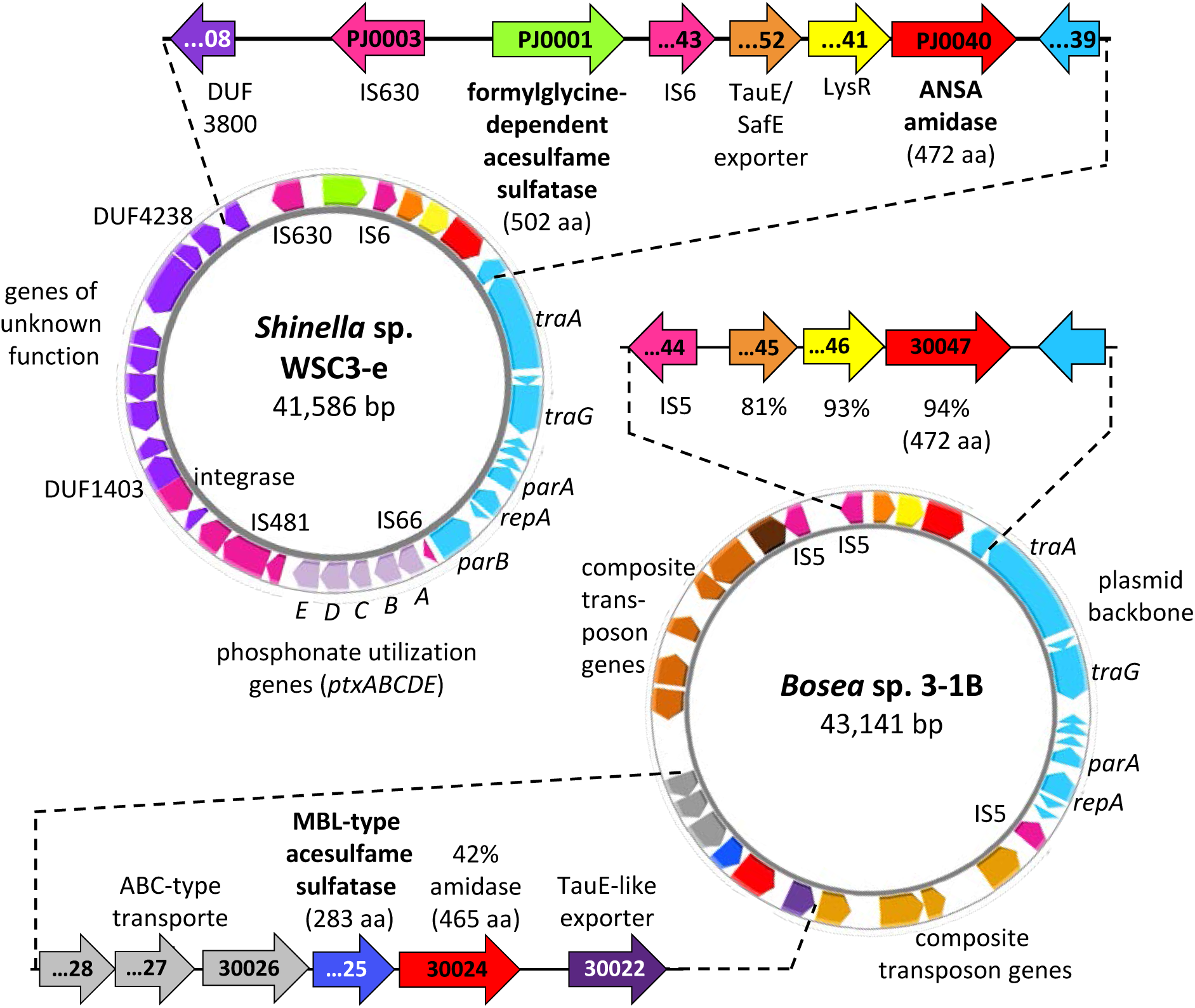
Comparison of AUP plasmids between acesulfame-degrading *Shinella* and *Bosea*/*Chelatococcus* strains. In *Bosea* sp. 3-1B,^30^ an MBL-type acesulfame sulfatase is encoded in a six coding sequences spanning metabolic gene cluster embedded in a composite transposon, while the ANSA amidase gene is located in another gene cluster together with two coding sequences encoding a LysR-type regulator and a TauE/SafE export system. This ANSA amidase cluster is fully conserved in *Shinella* sp. WSC3-e. However, the MBL-type acesulfame sulfatase is replaced by a formylglycine-dependent acesulfame sulfatase, which is not encoded in a separate gene cluster but directly upstream of the ANSA amidase cluster. Labels within the gene symbols refer to locus tags (prefix BOSEA31B_ and SHIWSC3_ for strains 3-1B and WSC3-e, respectively). The amino acid sequence identity (blastp) of *Bosea* ANSA amidase, LysR and TauE/SafE proteins with the respective *Shinella* sequences is indicated. Additionally, the identity (blastp) of the second *Bosea* AUP plasmid amidase (BOSEA31B_30024) with the *Shinella* ANSA amidase is given. Both AUP plasmids are shown true to scale. Annotation details for the *Shinella* AUP plasmid are given in Table S3. The figure was generated with Proksee.^35^

On the other hand, the *Shinella* AUP plasmid harbors a gene cluster annotated to encode proteins involved in phosphonate utilization, several insertion sequence (IS) elements and other genes that are not present on the *Bosea* and *Chelatococcus* AUP plasmids (Figure 1, Table S3). One of these genes (SHIWSC3_PJ0001) is located directly upstream of the ANSA amidase gene cluster and encodes a protein annotated as formylglycine-dependent sulfatase with a length of 502 aa. Although the sulfatase motif (CxPxRxxxLTGR) essential for the posttranslational modification of a reactive site cysteine residue (C64 in the *Shinella* sulfatase) into the catalytically active formylglycine residue^37^ is conserved, the enzyme is only distantly related to characterized formylglycine-dependent sulfatases. For example, the *Shinella* sulfatase has 30% amino acid identity along 93% of the query sequence with the choline sulfatase from *Ensifer meliloti* (UniProt ID: O69787). Therefore, its function and substrate specificity cannot be clearly assigned by sequence comparison. However, considering the absence of the *Bosea*/*Chelatococcus* MBL-type acesulfame sulfatase gene cluster in the *Shinella* strains^30^ and the necessity of a sulfatase to catalyze acesulfame degradation, we hypothesize that the SHIWSC3_PJ0001 gene product encodes a novel acesulfame sulfatase.

### Mn^2+^-independent hydrolysis of acesulfame in *Shinella* sp. WSC3-e

For heterologous expression, we then cloned the sulfatase gene SHIWSC3_PJ0001 into *E. coli* Lemo21 (DE3). In line with our assumption that the *Shinella* gene encodes an acesulfame sulfatase, crude extracts of the transformed *E. coli* strain stoichiometrically converted acesulfame to ANSA, while a negative control obtained from cells incubated under the same conditions without the *Shinella* sulfatase gene did not (Figure 2A). This result also indicates that, in contrast to the *Bosea*/*Chelatococcus* MBL-type acesulfame sulfatase, the SHIWSC3_PJ0001-encoded enzyme is Mn^2+^-independent, as the lysogeny broth medium used for the heterologous expression in *E. coli* contains insufficient amounts of Mn^2+^ to enable appropriate loading of heterologous Mn^2+^-dependent proteins with this metal ion.^38^ Such Mn^2+^- independence of the *Shinella* acesulfame sulfatase was also corroborated by growing *Shinella* sp. WSC3-e on acesulfame as the sole carbon and energy source in a mineral salt medium that was either poor (0.15 µM) or rich (3.85 µM) in Mn^2+^, resulting in almost identical acesulfame transformation activity and growth (Figure 2B). In comparison, it was previously demonstrated that only the Mn^2+^-rich but not the Mn^2+^-poor mineral salt medium supports efficient acesulfame degradation of *Bosea* sp. 100-5.^30^

**Figure 2.**
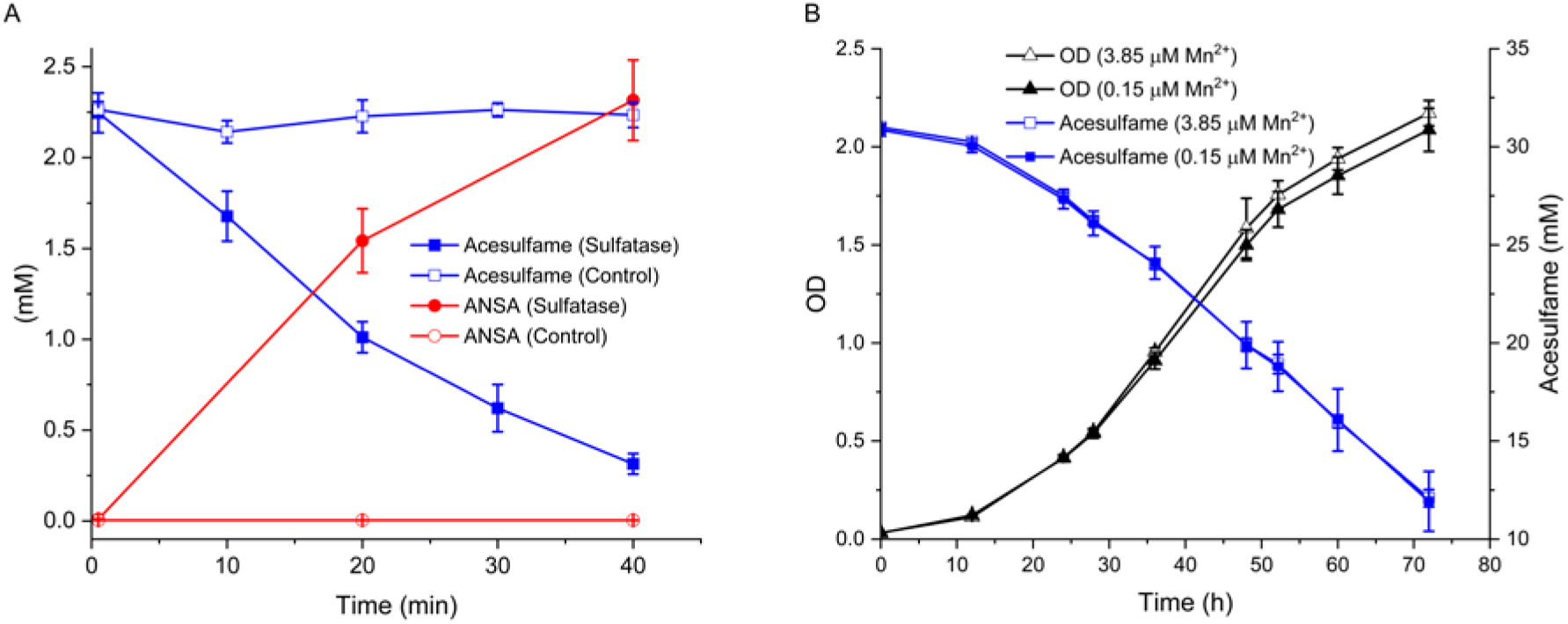
Biochemical and physiological characteristics of acesulfame degradation in *Shinella* sp. WSC3-e. (A) Stoichiometric conversion of acesulfame to ANSA by crude extracts from *E. coli* Lemo21 (DE3) cells expressing the *Shinella* sp. WSC3-e sulfatase gene SHIWSC3_PJ0001 (assay with 0.58 µg total protein μL^-1^). The corresponding crude extract of an *E. coli* Lemo21 (DE3) strain without the SHIWSC3_PJ0001 gene was used as a control (0.57 µg total protein μL^-1^) and did neither show acesulfame removal nor ANSA formation (detection limit: 0.2 μM with LC-MS/MS). (B) Incubation of *Shinella* sp. WSC3-e on acesulfame at 30°C in Mn^2+^-poor mineral salt medium (0.15 µM Mn^2+^) with or without Mn^2+^ supplementation (+3.7 µM) enabled growth with a doubling time of about 7 hours. OD, optical density at 600 nm. Values given represent mean and standard deviation from five biological replicates.

### Proteomic analysis of *Shinella* sp. WSC3-e

Proteomic analysis (three replicates) of the crude extract of the acesulfame-grown *Shinella* sp. WSC3-e identified 17,839 peptides grouping into 2,498 proteins, out of 8,823 proteins predicted from the genome. The *Shinella* AUP plasmid-encoded SHIWSC3_PJ0001 sulfatase and the SHIWSC3_PJ0040 amidase were found among the 20 most abundant proteins (Table 1, Table S4), accounting for 6.7% and 1.2% of the proteome, respectively. Besides, several chromosomal genes encoding ABC-type transporter proteins and major chaperons (GroEL/GroES, DnaK systems) as well as a plasmid-borne (plasmid PD, ENA/GenBank ID: OZ000534) gene encoding a chaperon ClpB-related protein were also strongly expressed. In regard to basic cellular functions, chromosome-encoded elongation factors and enzymes for the tricarboxylic acid cycle, for ammonia assimilation and for the glyoxylate cycle were most prominent. Additionally, several genes from another plasmid (plasmid PF, ENA/GenBank ID: OZ000536) were highly expressed (Table 1). These encode an acetyl-CoA C transferase (SHIWSC3_PF0360 gene product) and a couple of putative enzymes with less reliable functional annotation.

**Table 1.**
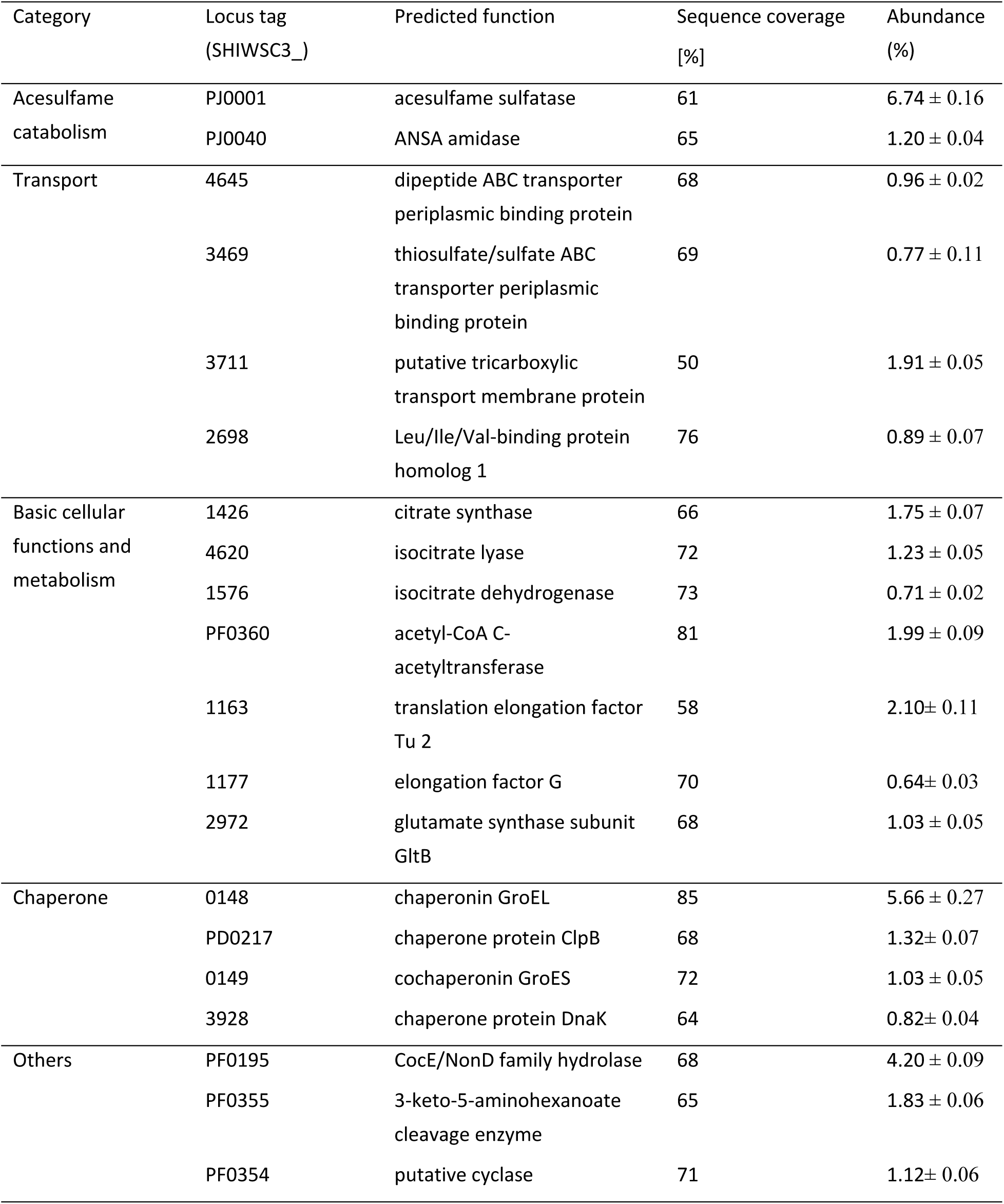
The 20 most abundant proteins in *Shinella* sp. WSC3-e

| Category | Locus tag<br>(SHIWSC3_) | Predicted function | Sequence coverage<br>[%] | Abundance<br>(%) |
| --- | --- | --- | --- | --- |
| Acesulfame<br>catabolism | PJ0001 | acesulfame sulfatase | 61 | 6.74 ± 0.16 |
|  | PJ0040 | ANSA amidase | 65 | 1.20 ± 0.04 |
| Transport | 4645 | dipeptide ABC transporter<br>periplasmic binding protein | 68 | 0.96 ± 0.02 |
|  | 3469 | thiosulfate/sulfate ABC<br>transporter periplasmic<br>binding protein | 69 | 0.77 ± 0.11 |
|  | 3711 | putative tricarboxylic<br>transport membrane protein | 50 | 1.91 ± 0.05 |
|  | 2698 | Leu/Ile/Val-binding protein<br>homolog 1 | 76 | 0.89 ± 0.07 |
| Basic cellular<br>functions and<br>metabolism | 1426 | citrate synthase | 66 | 1.75 ± 0.07 |
|  | 4620 | isocitrate lyase | 72 | 1.23 ± 0.05 |
|  | 1576 | isocitrate dehydrogenase | 73 | 0.71 ± 0.02 |
|  | PF0360 | acetyl-CoA C-acetyltransferase | 81 | 1.99 ± 0.09 |
|  | 1163 | translation elongation factor<br>Tu 2 | 58 | 2.10± 0.11 |
|  | 1177 | elongation factor G | 70 | 0.64± 0.03 |
|  | 2972 | glutamate synthase subunit<br>GltB | 68 | 1.03 ± 0.05 |
| Chaperone | 0148 | chaperonin GroEL | 85 | 5.66 ± 0.27 |
|  | PD0217 | chaperone protein ClpB | 68 | 1.32± 0.07 |
|  | 0149 | cochaperonin GroES | 72 | 1.03 ± 0.05 |
|  | 3928 | chaperone protein DnaK | 64 | 0.82± 0.04 |
| Others | PF0195 | CocE/NonD family hydrolase | 68 | 4.20 ± 0.09 |
|  | PF0355 | 3-keto-5-aminohexanoate<br>cleavage enzyme | 65 | 1.83 ± 0.06 |
|  | PF0354 | putative cyclase | 71 | 1.12± 0.06 |

Protein abundance and acesulfame sulfatase activity were analyzed in SEC fractions obtained from crude extracts of acesulfame-grown *Shinella* sp. WSC3-e. Acesulfame degradation activity peaked in fraction F10 (Figure 3A), corresponding to an apparent molecular mass of about 200 kDa. The SHIWSC3_PJ0001 sulfatase had its highest abundance in fraction F10 too, indicating its presence as a homo-trimeric (171 kDa) or homo-tetrameric (228 kDa) form, based on a predicted monomeric size of 57 kDa. In addition, several other proteins present at lower abundance than the sulfatase co-eluted with acesulfame degradation (Figure 3A). In a more comprehensive hierarchical cluster analysis, the acesulfame hydrolyzing activity was analyzed together with the distribution of the 200 most abundant proteins found in the SEC fractions (Figure S2, Table S5). Five proteins showed a strong correlation between their abundance distribution and the distribution of acesulfame sulfatase activity across the SEC fractions (Figure 3B). Among these five proteins, the SHIWSC3_PJ0001 sulfatase was the only hydrolase according to the annotation.

**Figure 3.**
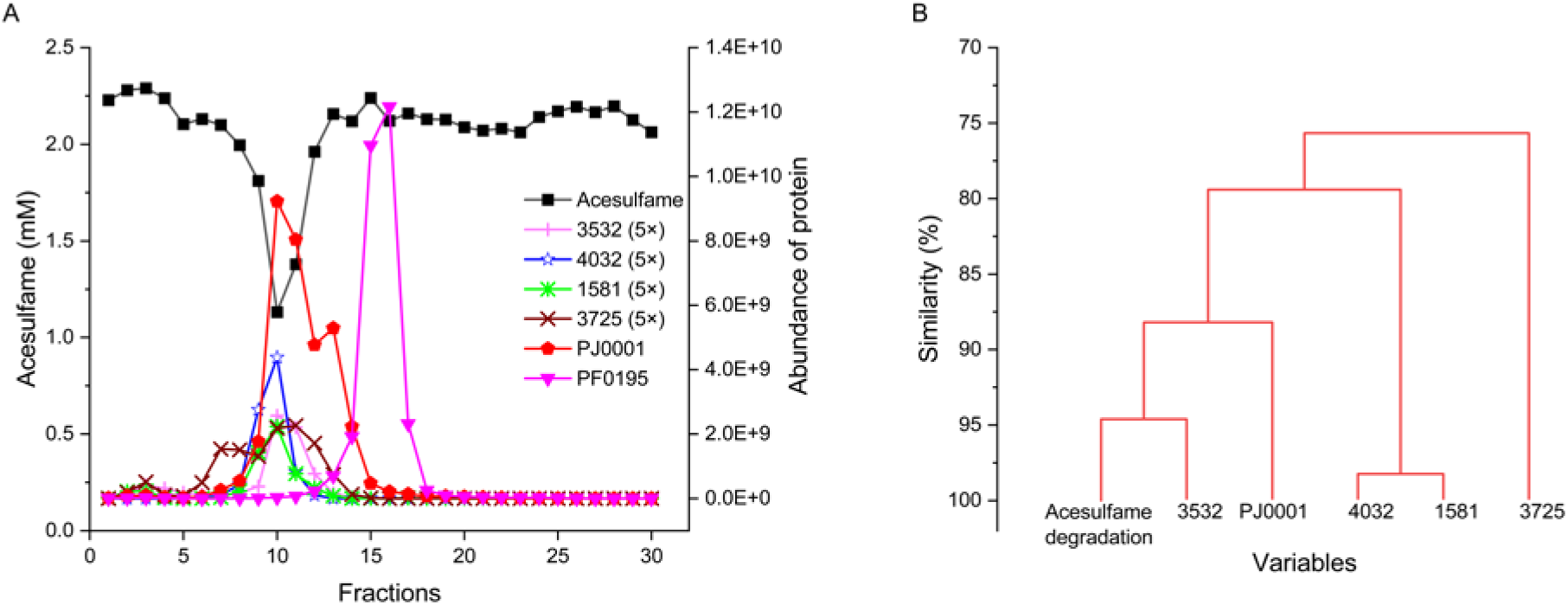
Fractionation of *Shinella sp.* strain WSC3-e crude extract by SEC and correlation of protein abundance with acesulfame sulfatase activity in the fractions. Locus tag prefix of strain WSC3-e proteins is SHIWSC3_. (A) SEC elution profiles of acesulfame degradation (remaining acesulfame concentration after incubating a subsample of each fraction with 2.25 mM acesulfame for 40 min) and abundance of the SHIWSC3_PJ0001 sulfatase, several co-eluting proteins and the SHIWSC3_PF0195 CocE/NonD family hydrolase. For better visibility, abundance values of proteins that co-elute with acesulfame degradation activity were five-fold increased (5x), except SHIWSC3_PJ0001. (B) Sub-dendrogram of the hierarchical cluster analysis for acesulfame hydrolysis and abundance distribution of the 200 most abundant proteins found in the SEC fractions. Only the five proteins with a similarity value >75% are shown in this sub-dendrogram.

### Acesulfame sulfatase and ANSA amidase genes in public sequence databases

As outlined above, the *Shinella* AUP plasmid bearing the Mn^2+^-independent acesulfame sulfatase and the ANSA amidase genes is 100% conserved in all acesulfame-degrading *Shinella* strains isolated thus far. Certain features of the plasmid were also found to be 100% conserved in some other bacteria, e.g., the IS481 family element together with the phosphonate utilization gene cluster in the genomes of *Agrobacterium tumefaciens* CFBP6623 (GenBank ID: GCA_005221385.1), *Brucella tritici* strain SY-3 (GenBank ID: GCA_023108915.1) and *Ochrobactrum* sp. WY7 (GenBank ID: GCA_018437805.1). In contrast, for the two hydrolase genes involved in acesulfame degradation, such 100% sequence conservation could not be detected in other published genome datasets. For the Mn^2+^-independent acesulfame sulfatase gene, the best match showing 85% nucleotide identity (at 97% query coverage and 1% gaps in the alignment) is a gene from *Devosia oryzisoli* PTR5 encoding a 484-aa protein (NCBI ID: WP_191775306.1) that shares 91% identical amino acid residues with the *Shinella* enzyme. Other closely related sequences (>86% protein sequence identity) are present, e.g., in *Devosia nitrariae* NBRC 112416 (WP_284342494.1), *Rhizobium* sp. S152 (WP_289722500.1) and *Paradevosia shaoguanensis* M48 (WP_281736881.1). However, genomes of these strains do not possess a sequence with significant similarity (as defined by the NCBI megablast algorithm) to the *Shinella* ANSA amidase gene. Moreover, the latter gene shows high identity (92% at 100% query coverage) only with the gene encoding the ANSA amidase in the previously described acesulfame-degrading *Bosea* and *Chelatococcus* strains,^30^ resulting in 94% identity of the protein sequence.

In summary, even though orthologous sequences highly similar with the *Shinella* acesulfame sulfatase and ANSA amidase exist in the databases, identical gene sequences (≥97% identity) are not present in published genomes, metagenome assembled genomes (MAGs) or any other wastewater-associated metagenome assembly searched in this study. Therefore, we extended the blastn search for the *Shinella* acesulfame sulfatase and ANSA amidase genes to SRA files from shotgun metagenome and metatranscriptome sequencing projects related to wastewater, WWTP or receiving waters. In total, more than 5,300 SRA experiment files resulting from sampling campaigns between the years 2005 and 2023 and covering all populated continents were analyzed, amounting to 59.9 tera base pairs (Tbp) of nucleotide sequence data (Table 2, Table S1). In order to compare the occurrence of both the Mn^2+^- dependent and the Mn^2+^-independent acesulfame degradation pathways, we also searched for the *Bosea*/*Chelatococcus* acesulfame sulfatase and ANSA amidase genes. As expected from the previous detection of the *Bosea* sp. 100-5 AUP plasmid in several metagenome assemblies,^30^ the corresponding acesulfame sulfatase and ANSA amidase genes were regularly found in SRA files from most geographical regions (Table 2, Figure 4A), albeit often at low coverage that did not allow assembly of complete gene sequences (Figure S3). The earliest datasets with the *Bosea*/*Chelatococcus* amidase gene were from 2012 (Sweden, PRJEB14051, and Hong Kong, PRJNA432264) and with the *Bosea*/*Chelatococcus* sulfatase genes from 2013 (Argentina, PRJNA288131, USA, PRJNA286671 and PRJNA236782, Hong Kong, PRJNA478263, and Luxembourg, PRJNA230567). Both hydrolase genes are almost absent only in samples from Africa and South Asia, which are highly underrepresented in the SRA. Overall, the number of worldwide sampling sites possessing the *Bosea*/*Chelatococcus* acesulfame pathway correlated with the searched SRA dataset size (Figure 4B), i.e., the acesulfame sulfatase and ANSA amidase genes were detected approximately once every 260 and 220 giga base pairs (Gbp) of SRA experiments searched, respectively. In stark contrast to the abundance of the *Bosea*/*Chelatococcus* sequences, the *Shinella* acesulfame sulfatase and ANSA amidase genes were rare in the SRA files (Table 2) and could only be found in a couple of datasets resulting from sampling campaigns in China, Mexico, Finland and Hungary within the years 2020 to 2022 (Table 3).

**Figure 4.**
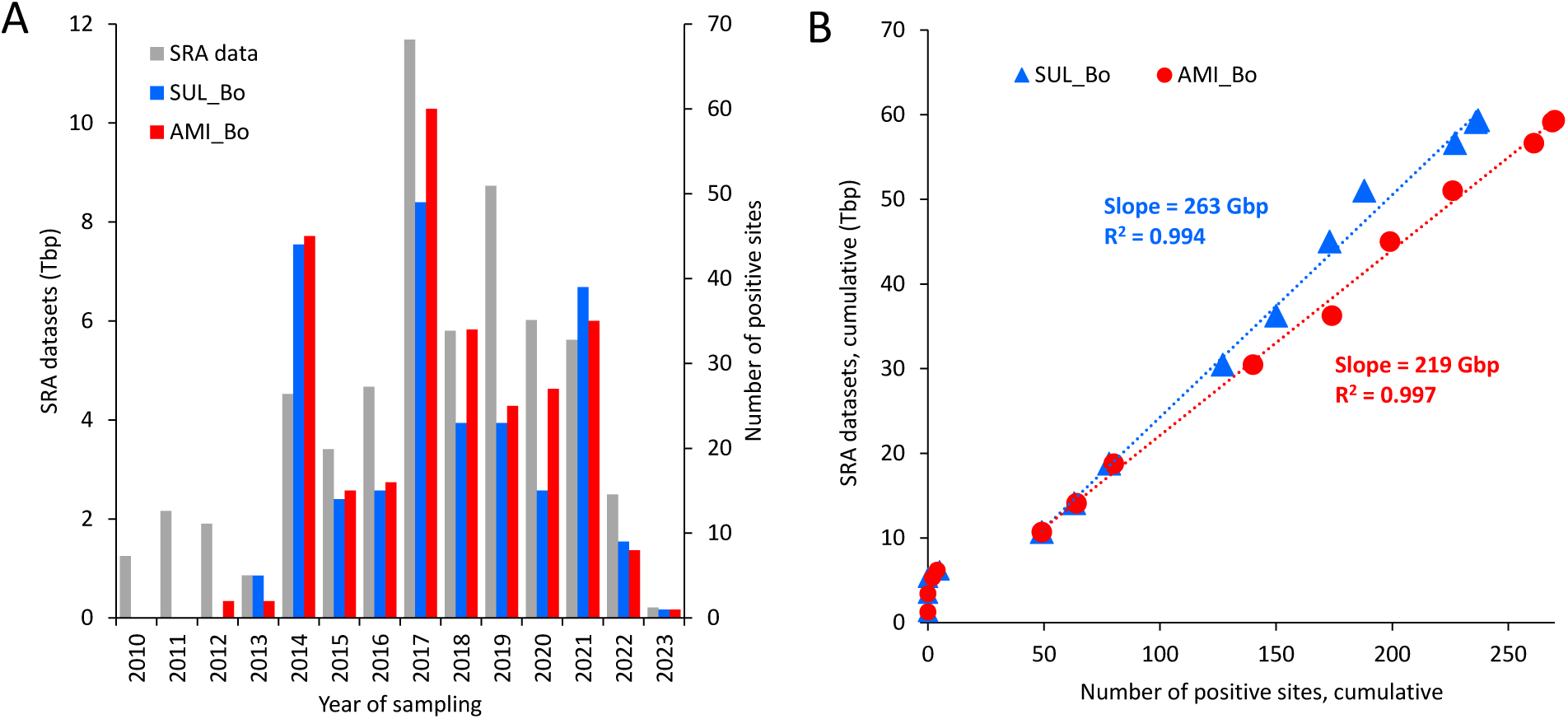
Historic occurrence of the *Bosea*/*Chelatococcus* acesulfame sulfatase (SUL_Bo) and ANSA amidase (AMI_Bo) genes in shotgun metagenome and metatranscriptome SRA datasets related to wastewater, WWTP or receiving waters across all populated continents. (A) Size of datasets searched by the blastn tool for the sulfatase and amidase genes as well as the number of positive sites considering sampling campaigns from 2010 to 2023. (B) Corresponding cumulative plot of SRA datasets searched versus the number of positive sampling sites. From the sampling year 2014 on, a linear correlation between SRA dataset size and number of positive sites turned up for both genes, for which the slopes and R^2^ values are given.

**Table 2.** Geographic distribution and number of sampling sites in which we detected nucleotide sequences identical (≥97%) to the acesulfame sulfatase (SUL_) and ANSA amidase (AMI_) genes from the acesulfame-degrading *Bosea*/*Chelatococcus* (Bo) and *Shinella* (Sh) strains in SRA datasets related to wastewater, WWTP or receiving waters. A complete list of all shotgun metagenome and metatranscriptome projects retrieved for the blastn search can be found in Table S1.

| Continent | Datasets (Tbp) | Campaign period | Sampling sites | Sites with detection of |  |  |  |
| --- | --- | --- | --- | --- | --- | --- | --- |
|  |  |  |  | SUL_Bo | AMI_Bo | SUL_Sh | AMI_Sh |
| North America | 11.4 | 2005–2022 | 277 | 50 | 62 | 2 | 2 |
| South America | 1.4 | 2008–2019 | 100 | 7 | 7 | 0 | 0 |
| Europe | 16.7 | 2010–2022 | 428 | 81 | 98 | 2 | 1 |
| Africa | 2.6 | 2013–2019 | 105 | 1 | 0 | 0 | 0 |
| West/Central Asia | 3.8 | 2015–2020 | 46 | 6 | 4 | 0 | 0 |
| East Asia | 18.6 | 2007–2023 | 290 | 84 | 90 | 3 | 3 |
| Southeast Asia | 3.8 | 2010–2020 | 60 | 5 | 5 | 0 | 0 |
| South Asia | 0.8 | 2013–2022 | 39 | 0 | 0 | 0 | 0 |
| Oceania | 0.9 | 2008–2019 | 24 | 3 | 4 | 0 | 0 |

**Table 3.** SRA datasets of metagenome projects in which we detected nucleotide sequences identical (≥97%) to the *Shinella* acesulfame sulfatase and ANSA amidase genes (abbreviations used for genes are as in Table 2).

| BioProject | Dataset (Gbp) | Country/region | Sampling sites | Year of sampling | Detected SUL and AMI genes |
| --- | --- | --- | --- | --- | --- |
| PRJNA807808<sup>39</sup> | 38.9 | **Mexico/**<br>Querétaro | domestic wastewater and pilot-scale WWTP | 2020 | SUL_Sh, AMI_Sh, SUL_Bo, AMI_Bo |
| PRJNA983699 | 24.1 | **Mexico/**<br>Querétaro | pilot-scale WWTP and anaerobic digester | 2021 | SUL_Sh, AMI_Sh, SUL_Bo |
| PRJNA876047 | 36.2 | **China/**<br>Shijiazhuang | hospital wastewater | 2020 | SUL_Sh, AMI_Sh, AMI_Bo |
| PRJNA876047 | 108.8 | **China/**<br>Shijiazhuang | hospital wastewater | 2021 | SUL_Sh, AMI_Sh, SUL_Bo, AMI_Bo |
| PRJNA988425 | 86.2 | **China/**<br>Xinjiang | WWTP | 2021 | SUL_Sh, AMI_Sh, AMI_Bo |
| PRJEB61958 | 160.7 | **Finland/**<br>Oulu | WWTP | 2021 | SUL_Sh |
| PRJEB61958 | 487.5 | **Finland/**<br>Oulu | WWTP | 2022 | SUL_Sh, AMI_Sh, SUL_Bo, AMI_Bo |
| PRJNA929705 | 66.6 | **Hungary/**<br>Kecskemt and Szeged | anaerobic digester in WWTP and biogas plants | 2021 | AMI_Sh (Kecskemt biogas plant), SUL_Bo (Szeged WWTP) |

## Discussion

Recently, a Mn^2+^-dependent two-step hydrolysis pathway for the degradation of the artificial sweetener acesulfame has been elucidated in *Bosea* and *Chelatococcus* strains isolated from WWTPs located in Germany and Hong Kong.^29,30^ In the current study, we identified and characterized a variation of this pathway in *Shinella* strains isolated from wastewater in which the first step of acesulfame hydrolysis is catalyzed by a novel Mn^2+^-independent, formylglycine-dependent sulfatase instead of the Mn^2+^- dependent MBL-type sulfatase in *Bosea*/*Chelatococcus*. The subsequent step for the hydrolysis of ANSA to the common metabolite acetoacetate is catalyzed by an amidase signature family enzyme conserved in both pathways.

The enzymes of the *Bosea*/*Chelatococcus* as well as the *Shinella* acesulfame degradation pathway are encoded by genes located on small conjugative plasmids (AUP plasmids) of about 40 kbp in size. While some diversity among the AUP plasmids have been observed in *Bosea* and *Chelatococcus* strains, e.g., the loss of non-essential genes and rearrangements of larger plasmid segments,^30^ all acesulfame-degrading *Shinella* isolates possess a 100% conserved AUP plasmid. This high conservation might be indicative of a very recent recombination event that gave rise to the *Shinella* AUP plasmid. Although the formylglycine-dependent sulfatase has not been detected in other bacteria yet, its horizontal transfer to other wastewater populations seems possible, as the gene is independent from complex gene environments and already located in a mobile element. In comparison, the special requirement on Mn^2+^ uptake and selective loading of the MBL-type sulfatase with Mn^2+^ might restrict horizontal gene transfer of the *Bosea*/*Chelatococcus* acesulfame degradation pathway. In this context, it has been reported that species related to the genera *Deinococcus*, *Methylobacterium*, *Bosea* and *Chelatococcus* accumulate high intracellular Mn^2+^ concentrations.^40^

The biochemical function of the *Shinella* formylglycine-dependent acesulfame sulfatase in converting acesulfame to ANSA was proven by establishing the enzyme activity in *E. coli* heterologously expressing the *Shinella* gene (SHIWSC3_PJ0001 from *Shinella* sp. WSC3-e). Moreover, for acesulfame-grown *Shinella* sp. WSC3-e, proteomic analysis and activity assays on the SEC fractionated proteome confirmed the expression of the SHIWSC3_PJ0001 gene and the physiological role of its gene product in hydrolyzing acesulfame. Theoretically, other hydrolases present in the proteome might also contribute to acesulfame degradation *in vivo*. The SHIWSC3_PF0195 gene, e.g., is strongly expressed in acesulfame-grown *Shinella* sp. WSC3-e (Table 1) and encodes a CocE/NonD family hydrolase, predicted to catalyze cleavage of esters.^41^ However, acesulfame hydrolysis activity in SEC fractions was only associated with the SHIWSC3_PJ0001 gene product, but not with other hydrolases, indicating that the SHIWSC3_PJ0001 gene product is exclusively responsible for acesulfame hydrolysis *in vivo* (Figure 3). Proteomic analysis also confirmed the high-level expression of the ANSA amidase gene (SHIWSC3_PJ0040) in the *Shinella* strain, supporting its role in hydrolyzing ANSA *in vivo*.

Several transport protein candidates encoded in the genome of *Shinella* sp. WSC3-e may be involved in the uptake of the negatively charged acesulfame. One example is the ABC-type transporter encoded by loci SHIWSC3_PJ0026, _PJ0027 and _PJ0028 on the *Shinella* AUP plasmid. However, the encoded proteins were not detected in the proteome of *Shinella* sp. WSC3-e (Table S4). Alternatively, the highly abundant, chromosome-encoded ABC-type periplasmic binding proteins (encoded by SHIWSC3_4645 and SHIWSC3_3469) (Table 1) together with the corresponding membrane and ATP-binding subunits could be involved in acesulfame import.

After the two-step hydrolysis of acesulfame, the common metabolite acetoacetate is formed, which is likely metabolized via activation to acetoacetyl-CoA and subsequent cleavage to two acetyl-CoA.^28^ As indicated by the proteome of acesulfame-grown *Shinella* sp. WSC3-e (Table 1, Table S4), acetyl-CoA is then dissimilated via the tricarboxylic acid cycle, while assimilation proceeds via the glyoxylate cycle, with isocitrate lyase as the key enzyme. Due to the role of acetyl-CoA as central metabolite, other metabolic pathways might be employed as well, e.g., those present in strictly anaerobic microorganisms. Consequently, the acesulfame hydrolyzing pathway seems not to be restricted to the thus far isolated aerobic and facultative denitrifying bacterial strains. However, maturation of the *Shinella* acesulfame sulfatase is dependent on oxygen, as the essential posttranslational modification from the active site cysteine to a formylglycine residue is catalyzed by an oxygenase-like enzyme.^42^ Under the denitrifying conditions reported to enable acesulfame mineralization in *Shinella*,^31^ the required oxygen might be provided during nitrate reduction, as the conversion of the intermediate nitric oxide can lead to formation of molecular oxygen.^43^ In strictly anoxic environments, an alternative pathway would be necessary that employs a radical S-adenosylmethionine formylglycine-generating enzyme for the posttranslational modification of either an active site cysteine or a serine residue to formylglycine.^44,45^

Acesulfame degradation genes are only found at low frequencies in published metagenome and metatranscriptome assemblies related to wastewater environments. This might be due to the inherent limitation of the assembly algorithms: low abundance genes are often discarded by assemblers and, consequently, can be missing in MAGs or other assemblies.^46^ We therefore searched in the SRA database directly and found regular occurrence of the *Bosea*/*Chelatococcus* acesulfame sulfatase and ANSA amidase genes in WWTPs and related environments since 2013 (Figure 4, Table 2, Table S1).

The earliest shotgun metagenome and metatranscriptome studies dealing with full-scale WWTPs resulted from sampling campaigns around the years 2007 and 2009.^47-49^ Consequently, only a few datasets from this period could be retrieved. However, due to an increasing sequencing effort for wastewater-associated environments afterwards, the size of searched SRA sequences related to collection dates in 2010 and 2011 amounted already to 1.25 and 2.16 Tbp, respectively, and covered almost all continents (Table 2). Nevertheless, the acesulfame pathway could not be detected in these datasets, suggesting that the corresponding hydrolase genes were either introduced into wastewater habitats later or enriched over time. Interestingly, the emergence of *Bosea*/*Chelatococcus* sulfatase and amidase genes in the years 2012 / 2013 correlates well with the first detection of a substantial acesulfame removal in WWTPs.^23,50^ Such removal appears to be exclusively mediated by the *Bosea*/*Chelatococcus* enzymes, as the *Shinella* sulfatase and amidase genes could not be detected in samples earlier than 2020. The linear correlation between the number of positive sites for the *Bosea*/*Chelatococcus* genes and the size of the SRA datasets searched clearly indicates that the *Bosea*/*Chelatococcus* pathway is constantly present since its emergence (Figure 4B). This finding holds particularly true for wastewater habitats in North America, Europe and East Asia, as other geographical regions are underrepresented in the database (Table 2, Table S1). The observed frequency of the acesulfame pathway genes in the SRA database (Figure 4B) does not represent their real abundance in wastewater. Rather, it is heavily influenced by the average size of the sequencing projects, which depends on the number of samples taken per site and the sequencing depth. For example, the dataset size spanned from about 1 Gbp (e.g., PRJNA768945) to almost 600 Gbp (e.g., PRJNA226633) per SRA experiment file, which typically represents the DNA/RNA sequences of one sample.

As for *Shinella*, due to the low number of recently published datasets, the abundance and distribution of the *Shinella* pathway in wastewater habitats cannot be determined unambiguously. In most cases, SRA files related to metagenome and metatranscriptome projects are released with a delay of at least two years due to data processing and preparation of publications. Consequently, only a few datasets resulting from the sampling years 2020 / 2021 or later are currently available. Nevertheless, the detection in 2020 coincides with the first isolation of acesulfame-degrading *Shinella* strains from German WWTPs in the same year (*Shinella* sp. WSC3-e^30^ and WSD5-1).

Our approach to directly search the raw sequence data of shotgun metagenome and metatranscriptome projects may also be relevant for investigating the dispersal of other genetic markers, e.g., for antibiotics resistance. Accordingly, this search method has recently been applied in a related work for the detection of trimethoprim resistance in surface water and wastewater by analyzing 1.7 Tbp of SRA datasets.^51^ As demonstrated now for the *Shinella* hydrolase genes, rare sequences in metagenomes could be missed when only searching in assembled data. Despite their low abundance, these genes might still be of importance in the respective habitat or could become dominant upon enrichment. This has also been demonstrated by the exclusive enrichment and isolation of acesulfame-degrading *Shinella* strains from the Sha Tin WWTP in Hong Kong, while the acesulfame pathway genes could not be found in the corresponding MAGs.^31^

The analysis of metagenome data, the monitoring of acesulfame degradation activity in WWTPs^23,50^ and the isolation of acesulfame-degrading bacterial strains are consistent with an emergence of the Mn^2+^- dependent *Bosea*/*Chelatococcus* acesulfame degradation pathway not before the year 2012. Already after 2013, however, the respective sulfatase and amidase genes can be regularly detected in samples from wastewater treatment systems around the world, indicating a fast and unhindered spreading of the pathway. In line with this, the Mn^2+^-dependence of the pathway appears not to be a limitation, as manganese is a relatively abundant heavy metal in municipal wastewater due to various anthropogenic sources, such as ubiquitous steel abrasion and corrosion.^52^ Therefore, the emergence of an alternative Mn^2+^-independent acesulfame sulfatase eight years later than the *Bosea*/*Chelatococcus* enzyme, is somewhat surprising. Interestingly, both pathways may even coexist, as shown for sites in Mexico, Finland and China (Table 3). However, a temporary Mn^2+^ depletion in WWTPs, e.g., due to an microalgal bloom^53^ or when treating wastewater with microalgae as in project PRJNA807808^39^ (Table 2), might have facilitated the evolution and maintenance of the Mn^2+^-independent pathway, currently found exclusively in *Shinella* strains.

In conclusion, at least two different hydrolase mechanisms for acesulfame exist, which is impressive, as the catalyzed nucleophilic attack towards the sulfur atom is challenging due to acesulfame’s electron-rich ring system.^54^ Both the Mn^2+^-dependent and the Mn^2+^-independent sulfatases showed nature’s potential to provide efficient catalysts for chemically very challenging reactions.

## Acknowledgements

The authors thank Ute Lohse (UFZ) for cultivation of the strains, Benjamin Scheer (UFZ) for technical support in proteomic analysis, and Nadine Hellmold (UFZ) for establishing molecular mass standards for SEC calibration.

Y. L. was supported by the China Scholarship Council under the doctoral fellowship (202004910433).

Protein mass spectrometry was done at the Centre for Chemical Microscopy (ProVIS) at UFZ, which is supported by European regional development funds (EFRE—Europe Funds Saxony) and the Helmholtz Association. Data retrieval and local BLAST analysis of SRA datasets were performed at the High-Performance Computing Cluster EVE, a joint effort of the UFZ (http://www.ufz.de/) and the German Centre for Integrative Biodiversity Research (iDiv) Halle-Jena-Leipzig (http://www.idiv-biodiversity.de/).

## Supporting Information

Table S1. Searched shotgun metagenome/metatranscriptome projects related to wastewater, wastewater treatment and wastewater receiving waters.

Table S2. Genome comparison of acesulfame-degrading strains *Shinella* sp. WSC3-e, WSD5-1 and YE25.

Table S3. Coding sequences of the AUP plasmid from *Shinella* sp. WSC3-e with predicted functions.

Table S4. Identified proteins in the crude extract proteome of *Shinella* sp. WSC3-e.

Table S5. The 200 most abundant proteins found in SEC fractions.

Figure S1. Alignment of the *Shinella* sp. YE25 genome against the *Shinella* sp. WSC3-e genome.

Figure S2. Dendrogram of proteins for clustering analysis.

Figure S3. Examples for blastn results with acesulfame sulfatase genes.

---

**Figure S1.**
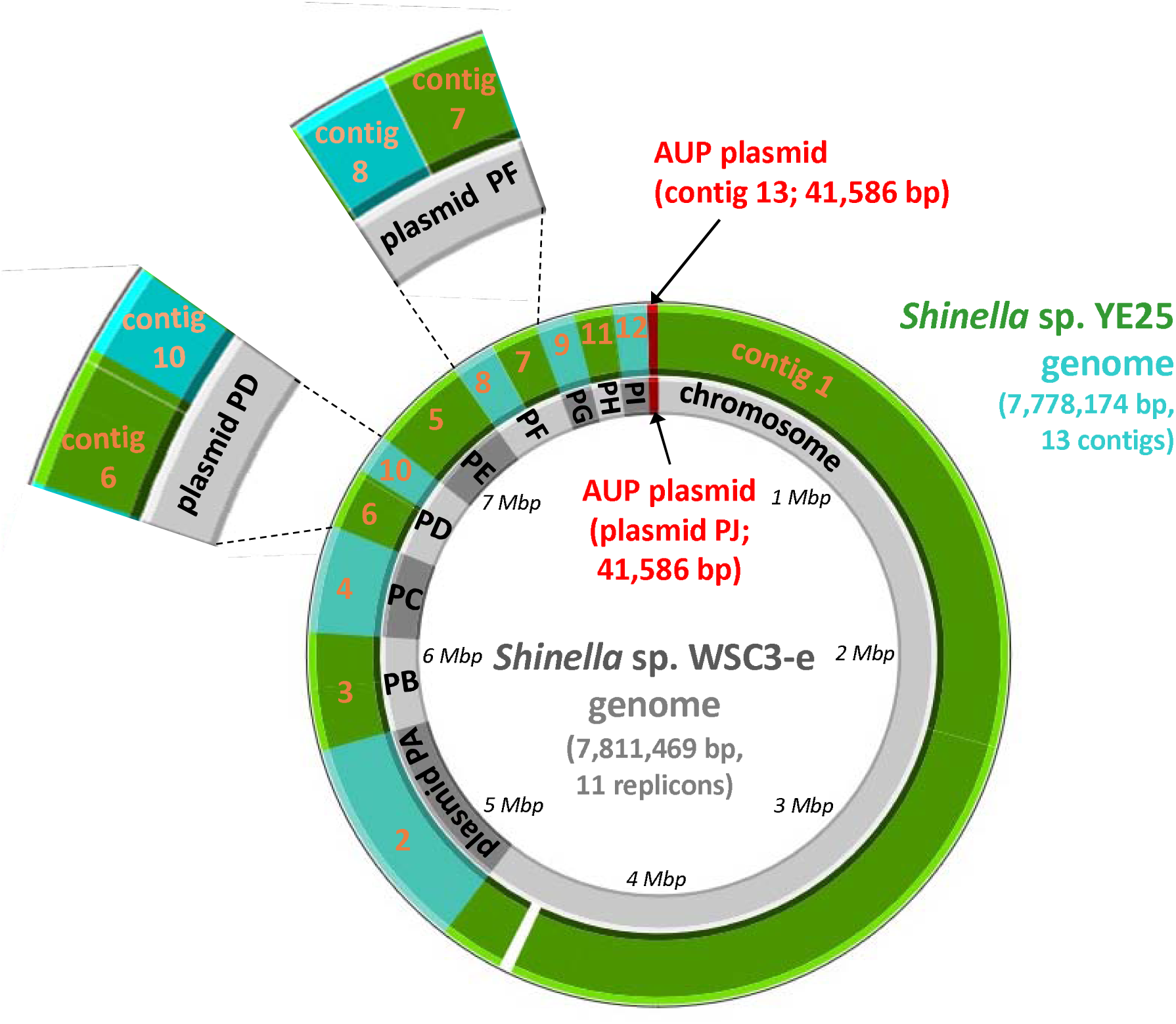
Alignment of the Shinella sp. YE25 genome (green and turquoise) against the Shinella sp. WSC3-e genome (gray). Respective contig numbers/replicon names are indicated (orange and black). Both strains share almost identical genome sequence (≥99%) and replicon organization, suggesting the presence of a large chromosome (about 4.7 Mbp) and several plasmids of various size. The smallest plasmid, for example, is the AUP plasmid (41,586 bp, highlighted in red) which is 100% identical in both strains. In contrast, a section (ca. 30 kbp) close to the end of the WSC3-e chromosome is completely missing in strain YE25. In addition, plasmids PD and PF of strain WSC3-e can be derived from smaller YE25 plasmids. In each case, two plasmids were rearranged to a larger one (see enlargement). The figure was generated with Proksee.

**Figure S2.**
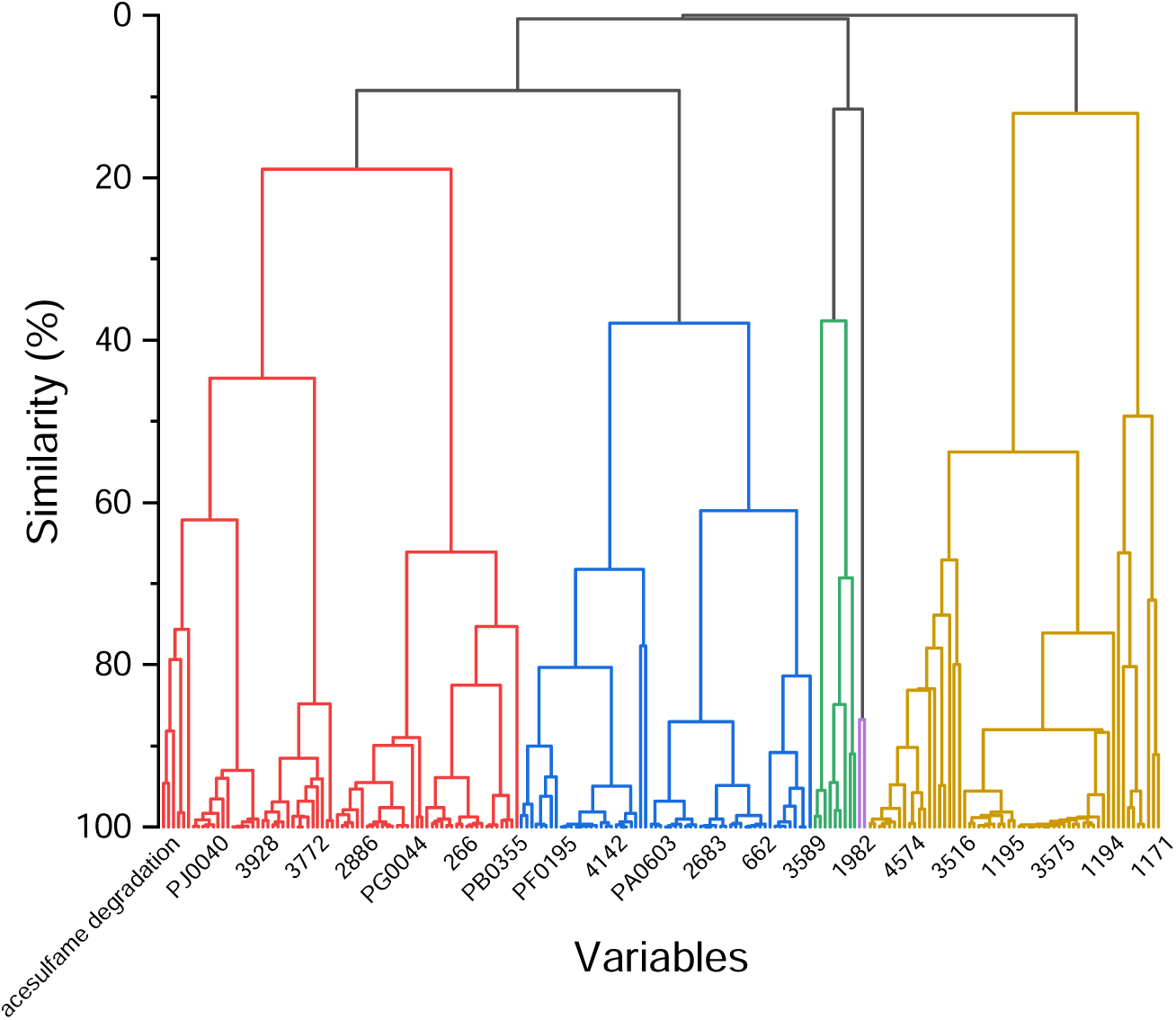
Dendrogram of hierarchical cluster analysis for acesulfame degradation and abundance distribution of 200 most abundant Shinella sp. WSC3-e proteins after summing up the proteins in 30 SEC fractions. Protein names indicated refer to locus tags (prefix SHIWSC3_). Not all protein names are shown due to space limitation. The protein names shown were automatically decided by the Origin software.

**Figure S3.**
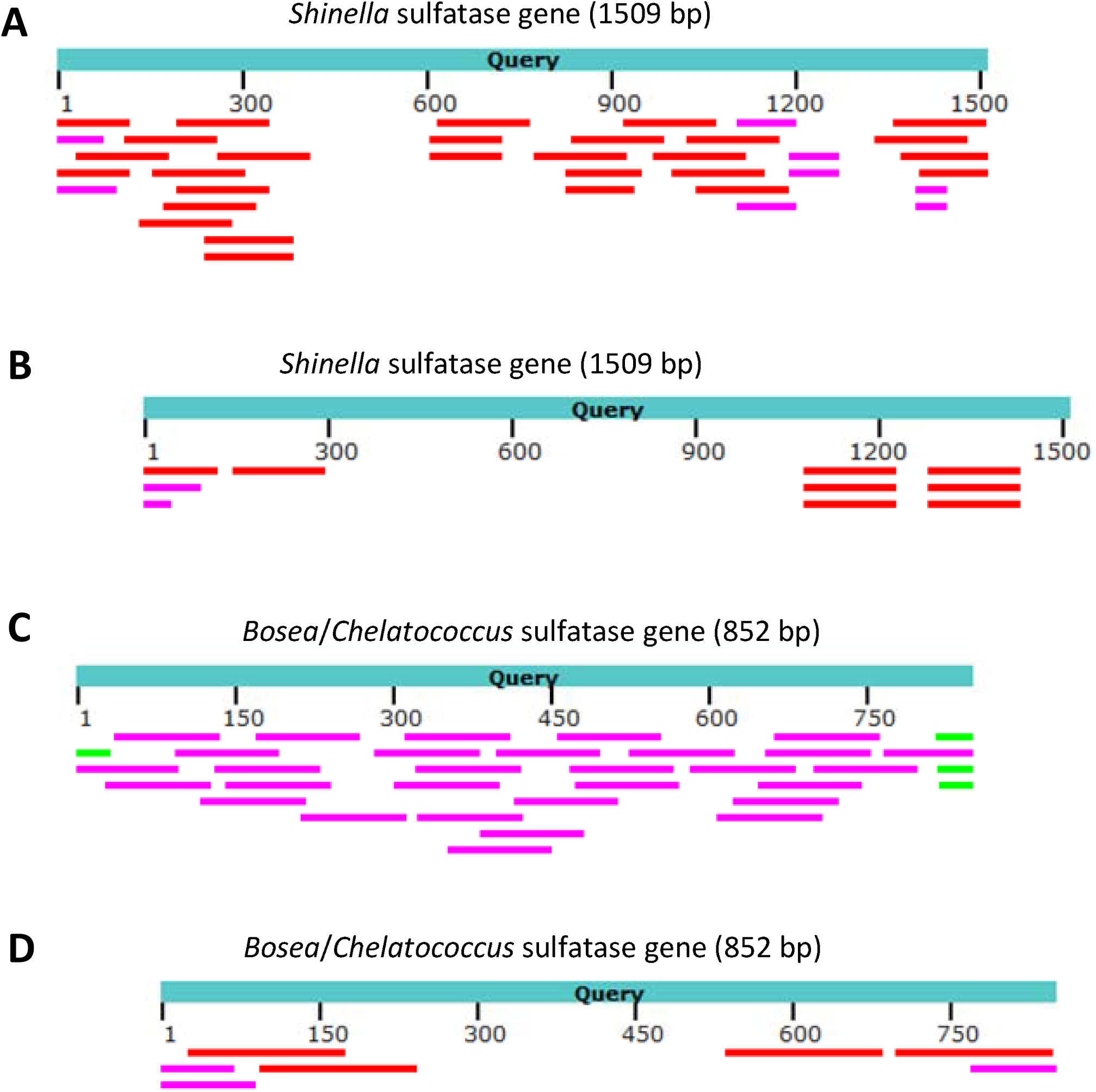
Examples for blastn results with acesulfame sulfatase genes from acesulfame-degrading Bosea/Chelatococcus and Shinella strains as query against datasets from the SRA database. For the alignments, only sequences showing ≥97% identity to the query sequence were considered. Color code as used by the NCBI blast graphic summary showing alignment scores (≥200, red; 80 – 200, magenta; 50 – 80, green). (A) Search with SHIWSC3_PJ0001 against SRA files from project PRJNA807808 (150-bp reads; pilot-scale microalgae-bacteria wastewater treatment system, Querétaro, Mexico, sampled in 2020). (B) Search with SHIWSC3_PJ0001 against SRA files from project PRJNA988425 (150-bp reads; WWTP in Xinjiang, China, sampled in 2021). (C) Search with BOSEA1005_40015 against SRA files from project PRJNA967004 (100-bp reads; WWTP in Galway, Ireland, sampled in 2020 and 2021). (D) Search with BOSEA1005_40015 against SRA files from project PRJNA952735 (150-bp reads; WWTP in Regina, Canada, sampled in 2017).

## References

(1) Roberts, M. W.; Wright, J. T., Nonnutritive, low caloric substitutes for food sugars: clinical implications for addressing the incidence of dental caries and overweight/obesity. Int. J. Dent. 2012, 2012, 625701.

(2) Klug, C.; von Rymon Lipinski, G.-W., Acesulfame K. In Sweeteners and Sugar Alternatives in Food Technology, 2012; pp 91–115.

(3) Lange, F. T.; Scheurer, M.; Brauch, H. J., Artificial sweeteners--a recently recognized class of emerging environmental contaminants: a review. Anal. Bioanal. Chem. 2012, 403 (9), 2503–18.

(4) Naik, A. Q.; Zafar, T.; Shrivastava, V. K., Environmental impact of the presence, distribution, and use of artificial sweeteners as emerging sources of pollution. J Environ Public Health 2021, 2021, 6624569.

(5) Gan, Z.; Sun, H.; Feng, B.; Wang, R.; Zhang, Y., Occurrence of seven artificial sweeteners in the aquatic environment and precipitation of Tianjin, China. Water Res. 2013, 47 (14), 4928–4937.

(6) Buerge, I. J.; Buser, H.-R.; Kahle, M.; Müller, M. D.; Poiger, T., Ubiquitous occurrence of the artificial sweetener acesulfame in the aquatic environment: an ideal chemical marker of domestic wastewater in groundwater. Environ. Sci. Technol. 2009, 43 (12), 4381–4385.

(7) Loos, R.; Carvalho, R.; António, D. C.; Comero, S.; Locoro, G.; Tavazzi, S.; Paracchini, B.; Ghiani, M.; Lettieri, T.; Blaha, L.; Jarosova, B.; Voorspoels, S.; Servaes, K.; Haglund, P.; Fick, J.; Lindberg, R. H.; Schwesig, D.; Gawlik, B. M., EU-wide monitoring survey on emerging polar organic contaminants in wastewater treatment plant effluents. Water Res. 2013, 47 (17), 6475–6487.

(8) Danner, L.; Malard, F.; Valdes, R.; Olivier-Van Stichelen, S., Non-nutritive sweeteners acesulfame potassium and sucralose are competitive inhibitors of the human p-glycoprotein/multidrug resistance protein 1 (PGP/MDR1). Nutrients. 2023, 15 (5), 1118.

(9) Chiang, Y.-F.; Chen, H.-Y.; Lai, Y.-H.; Ali, M.; Chen, Y.-C.; Hsia, S.-M., Consumption of artificial sweetener acesulfame potassium increases preterm risk and uterine contraction with calcium influx increased via myosin light chain kinase–myosin light chain 20 related signaling pathway. Mol. Nutr. Food Res. 2022, 66 (20), 2200298.

(10) Li, A. J.; Wu, P.; Law, J. C.-F.; Chow, C.-H.; Postigo, C.; Guo, Y.; Leung, K. S.-Y., Transformation of acesulfame in chlorination: Kinetics study, identification of byproducts, and toxicity assessment. Water Res. 2017, 117, 157–166.

(11) Ren, Y.; Geng, J.; Li, F.; Ren, H.; Ding, L.; Xu, K., The oxidative stress in the liver of *Carassius auratus* exposed to acesulfame and its UV irradiance products. Sci. Total Environ. 2016, 571, 755–762.

(12) Sang, Z.; Jiang, Y.; Tsoi, Y.-K.; Leung, K. S.-Y., Evaluating the environmental impact of artificial sweeteners: A study of their distributions, photodegradation and toxicities. Water Res. 2014, 52, 260–274.

(13) Li, A. J.; Schmitz, O. J.; Stephan, S.; Lenzen, C.; Yue, P. Y.-K.; Li, K.; Li, H.; Leung, K. S.-Y., Photocatalytic transformation of acesulfame: Transformation products identification and embryotoxicity study. Water Res. 2016, 89, 68–75.

(14) Bandyopadhyay, A.; Ghoshal, S.; Mukherjee, A., Genotoxicity testing of low-calorie sweeteners: aspartame, acesulfame-K, and saccharin. Drug Chem. Toxicol. 2008, 31 (4), 447–457.

(15) Hadidi, S.; Varmira, K.; Soltani, L., Evaluation of DNA damage induced by acesulfame potassium: spectroscopic, molecular modeling simulations and toxicity studies. J. Biomol. Struct. Dyn. 2022, 1–10.

(16) Colín-García, K.; Elizalde-Velázquez, G. A.; Gómez-Oliván, L. M.; García-Medina, S., Influence of sucralose, acesulfame-k, and their mixture on brain’s fish: A study of behavior, oxidative damage, and acetylcholinesterase activity in *Danio rerio*. Chemosphere. 2023, 340, 139928.

(17) Wiklund, A.-K. E.; Guo, X.; Gorokhova, E., Cardiotoxic and neurobehavioral effects of sucralose and acesulfame in *Daphnia*: Toward understanding ecological impacts of artificial sweeteners. Comp. Biochem. Physiol. C: Toxicol. Pharmacol. 2023, 273, 109733.

(18) Yu, Z.; Wang, Y.; Lu, J.; Bond, P. L.; Guo, J., Nonnutritive sweeteners can promote the dissemination of antibiotic resistance through conjugative gene transfer. ISME J. 2021, 15 (7), 2117–2130.

(19) Yu, Z.; Henderson, I. R.; Guo, J., Non-caloric artificial sweeteners modulate conjugative transfer of multi-drug resistance plasmid in the gut microbiota. Gut Microbes. 2023, 15 (1), 2157698.

(20) Yang, G.; Cao, J.-M.; Cui, H.-L.; Zhan, X.-M.; Duan, G.; Zhu, Y.-G., Artificial sweetener enhances the spread of antibiotic resistance genes during anaerobic digestion. Environ. Sci. Technol. 2023, 57 (30), 10919–10928.

(21) Roy, J. W.; Van Stempvoort, D. R.; Bickerton, G., Artificial sweeteners as potential tracers of municipal landfill leachate. Environ. Pollut. 2014, 184, 89–93.

(22) Zhao, Z.; Yin, H.; Xu, Z.; Peng, J.; Yu, Z., Pin-pointing groundwater infiltration into urban sewers using chemical tracer in conjunction with physically based optimization model. Water Res. 2020, 175, 115689.

(23) Castronovo, S.; Wick, A.; Scheurer, M.; Nödler, K.; Schulz, M.; Ternes, T. A., Biodegradation of the artificial sweetener acesulfame in biological wastewater treatment and sandfilters. Water Res. 2017, 110, 342–353.

(24) Kahl, S.; Kleinsteuber, S.; Nivala, J.; van Afferden, M.; Reemtsma, T., Emerging biodegradation of the previously persistent artificial sweetener acesulfame in biological wastewater treatment. Environ. Sci. Technol. 2018, 52 (5), 2717–2725.

(25) Burke, V.; Greskowiak, J.; Grünenbaum, N.; Massmann, G., Redox and temperature dependent attenuation of twenty organic micropollutants - a systematic column study. Water Environ. Res. 2017, 89 (2), 155–167.

(26) Marazuela, M. A.; Formentin, G.; Erlmeier, K.; Hofmann, T., Seasonal biodegradation of the artificial sweetener acesulfame enhances its use as a transient wastewater tracer. Water Res. 2023, 232, 119670.

(27) Marazuela, M. A.; Formentin, G.; Erlmeier, K.; Hofmann, T., Acesulfame allows the tracing of multiple sources of wastewater and riverbank filtration. Environ. Pollut. 2023, 323, 121223.

(28) Kleinsteuber, S.; Rohwerder, T.; Lohse, U.; Seiwert, B.; Reemtsma, T., Sated by a zero-calorie sweetener: wastewater bacteria can feed on acesulfame. Front. Microbiol. 2019, 10.

(29) Huang, Y.; Deng, Y.; Law, J. C.-F.; Yang, Y.; Ding, J.; Leung, K. S.-Y.; Zhang, T., Acesulfame aerobic biodegradation by enriched consortia and *Chelatococcus* spp.: Kinetics, transformation products, and genomic characterization. Water Res. 2021, 202, 117454.

(30) Bonatelli, M. L.; Rohwerder, T.; Popp, D.; Liu, Y.; Akay, C.; Schultz, C.; Liao, K.-P.; Ding, C.; Reemtsma, T.; Adrian, L.; Kleinsteuber, S., Recently evolved combination of unique sulfatase and amidase genes enables bacterial degradation of the wastewater micropollutant acesulfame worldwide. Front. Microbiol. 2023, 14.

(31) Huang, Y.; Yu, Z.; Liu, L.; Che, Y.; Zhang, T., Acesulfame anoxic biodegradation coupled to nitrate reduction by enriched consortia and isolated *Shinella* spp. Environ. Sci. Technol. 2022, 56 (18), 13096–13106.

(32) Kolmogorov, M.; Bickhart, D. M.; Behsaz, B.; Gurevich, A.; Rayko, M.; Shin, S. B.; Kuhn, K.; Yuan, J.; Polevikov, E.; Smith, T. P. L.; Pevzner, P. A., MetaFlye: scalable long-read metagenome assembly using repeat graphs. Nat. Methods. 2020, 17 (11), 1103–1110.

(33) Walker, B. J.; Abeel, T.; Shea, T.; Priest, M.; Abouelliel, A.; Sakthikumar, S.; Cuomo, C. A.; Zeng, Q.; Wortman, J.; Young, S. K.; Earl, A. M., Pilon: an integrated tool for comprehensive microbial variant detection and genome assembly improvement. PLOS ONE. 2014, 9 (11), e112963.

(34) Jain, C.; Rodriguez-R, L. M.; Phillippy, A. M.; Konstantinidis, K. T.; Aluru, S., High throughput ANI analysis of 90K prokaryotic genomes reveals clear species boundaries. Nat. Commun. 2018, 9 (1), 5114.

(35) Grant, J. R.; Enns, E.; Marinier, E.; Mandal, A.; Herman, E. K.; Chen, C.-y.; Graham, M.; Van Domselaar, G.; Stothard, P., Proksee: in-depth characterization and visualization of bacterial genomes. Nucleic Acids Res. 2023, 51 (W1), W484–W492.

(36) Ding, C.; Adrian, L., Comparative genomics in “*Candidatus* Kuenenia stuttgartiensis” reveal high genomic plasticity in the overall genome structure, CRISPR loci and surface proteins. BMC Genomics. 2020, 21 (1), 851.

(37) Carlson, B. L.; Ballister, E. R.; Skordalakes, E.; King, D. S.; Breidenbach, M. A.; Gilmore, S. A.; Berger, J. M.; Bertozzi, C. R., Function and structure of a prokaryotic formylglycine-generating enzyme*. J. Biol. Chem. 2008, 283 (29), 20117–20125.

(38) Anjem, A.; Varghese, S.; Imlay, J. A., Manganese import is a key element of the OxyR response to hydrogen peroxide in *Escherichia coli*. Mol. Microbiol. 2009, 72 (4), 844–58.

(39) Ovis-Sánchez, J. O.; Perera-Pérez, V. D.; Buitrón, G.; Quintela-Baluja, M.; Graham, D. W.; Morales-Espinosa, R.; Carrillo-Reyes, J., Exploring resistomes and microbiomes in pilot-scale microalgae-bacteria wastewater treatment systems for use in low-resource settings. Sci. Total Environ. 2023, 882, 163545.

(40) Fredrickson, J. K.; Li, S.-m. W.; Gaidamakova, E. K.; Matrosova, V. Y.; Zhai, M.; Sulloway, H. M.; Scholten, J. C.; Brown, M. G.; Balkwill, D. L.; Daly, M. J., Protein oxidation: key to bacterial desiccation resistance? ISME J. 2008, 2 (4), 393–403.

(41) Bresler Matthew, M.; Rosser Susan, J.; Basran, A.; Bruce Neil, C., Gene cloning and nucleotide sequencing and properties of a cocaine Esterase from *Rhodococcus* sp. Strain MB1. Appl. Environ. Microbiol. 2000, 66 (3), 904-908.

(42) Miarzlou, D. A.; Leisinger, F.; Joss, D.; Häussinger, D.; Seebeck, F. P., Structure of formylglycine-generating enzyme in complex with copper and a substrate reveals an acidic pocket for binding and activation of molecular oxygen. Chem. Sci. 2019, 10 (29), 7049–7058.

(43) Ettwig, K. F.; Butler, M. K.; Le Paslier, D.; Pelletier, E.; Mangenot, S.; Kuypers, M. M. M.; Schreiber, F.; Dutilh, B. E.; Zedelius, J.; de Beer, D.; Gloerich, J.; Wessels, H. J. C. T.; van Alen, T.; Luesken, F.; Wu, M. L.; van de Pas-Schoonen, K. T.; Op den Camp, H. J. M.; Janssen-Megens, E. M.; Francoijs, K.-J.; Stunnenberg, H.; Weissenbach, J.; Jetten, M. S. M.; Strous, M., Nitrite-driven anaerobic methane oxidation by oxygenic bacteria. Nature. 2010, 464 (7288), 543–548.

(44) Grove, T. L.; Lee, K.-H.; St. Clair, J.; Krebs, C.; Booker, S. J., In vitro characterization of AtsB, a radical SAM formylglycine-generating enzyme that contains three [4Fe-4S] clusters. Biochem. 2008, 47 (28), 7523–7538.

(45) Benjdia, A.; Leprince, J.; Guillot, A.; Vaudry, H.; Rabot, S.; Berteau, O., Anaerobic sulfatase-maturating enzymes: radical SAM enzymes able to catalyze in vitro sulfatase post-translational modification. J. Am. Chem. Soc. 2007, 129 (12), 3462–3463.

(46) Meziti, A.; Rodriguez, R. L.; Hatt, J. K.; Peña-Gonzalez, A.; Levy, K.; Konstantinidis, K. T., The reliability of metagenome-sssembled genomes (MAGs) in representing natural populations: snsights from comparing MAGs against isolate genomes derived from the same fecal sample. Appl. Environ. Microbiol. 2021, 87 (6).

(47) Sanapareddy, N.; Hamp Timothy, J.; Gonzalez Luis, C.; Hilger Helene, A.; Fodor Anthony, A.; Clinton Sandra, M., Molecular diversity of a north carolina wastewater treatment plant as revealed by pyrosequencing. Appl. Environ. Microbiol. 2009, 75 (6), 1688–1696.

(48) Albertsen, M.; Hansen, L. B. S.; Saunders, A. M.; Nielsen, P. H.; Nielsen, K. L., A metagenome of a full-scale microbial community carrying out enhanced biological phosphorus removal. ISME J. 2012, 6 (6), 1094–1106.

(49) Ibarbalz Federico, M.; Orellana, E.; Figuerola Eva, L. M.; Erijman, L., Shotgun metagenomic profiles have a high capacity to discriminate samples of activated sludge according to wastewater type. Appl. Environ. Microbiol. 2016, 82 (17), 5186–5196.

(50) Cardenas, M. A. R.; Ali, I.; Lai, F. Y.; Dawes, L.; Thier, R.; Rajapakse, J., Removal of micropollutants through a biological wastewater treatment plant in a subtropical climate, Queensland-Australia. J. environ. health sci. eng. 2016, 14 (1), 14.

(51) Kneis, D.; Lemay-St-Denis, C.; Cellier-Goetghebeur, S.; Elena, A. X.; Berendonk, T. U.; Pelletier, J. N.; Heß, S., Trimethoprim resistance in surface and wastewater is mediated by contrasting variants of the *dfrB* gene. ISME J. 2023, 17 (9), 1455–1466.

(52) Raclavska, H.; Drozdova, J.; Skrobankova, H.; Raclavsky, K., Behavior of metals in a combined wastewater collection system in Ostrava, Czech Republic. Water Environ. Res. 2015, 87 (2), 123–31.

(53) Vojvodić, S.; Dimitrijević, M.; Žižić, M.; Dučić, T.; Aquilanti, G.; Stanić, M.; Zechmann, B.; Danilović Luković, J.; Stanković, D.; Opačić, M.; Morina, A.; Pittman, J. K.; Spasojević, I., A three-step process of manganese acquisition and storage in the microalga *Chlorella sorokiniana*. J. Exp. Bot. 2023, 74 (3), 1107–1122.

(54) Popova, A. D.; Velcheva, E. A.; Stamboliyska, B. A., DFT and experimental study on the IR spectra and structure of acesulfame sweetener. J. Mol. Struct. 2012, 1009, 23–29.

